# Clonal evolution in serially passaged *Cryptococcus neoformans* x *deneoformans* hybrids reveals a heterogenous landscape of genomic change

**DOI:** 10.1101/2021.03.02.433606

**Authors:** Lucas A. Michelotti, Sheng Sun, Joseph Heitman, Timothy Y. James

## Abstract

*Cryptococcus neoformans* x *deneoformans* hybrids (also known as serotype AD hybrids) are basidiomycete yeasts that are common in a clinical setting. Like many hybrids, the AD hybrids are largely locked at the F1 stage and are mostly unable to undergo normal meiotic reproduction. However, these F1 hybrids, which display a high (∼10%) sequence divergence are known to genetically diversify through mitotic recombination and aneuploidy, and this diversification may be adaptive. In this study, we evolved a single AD hybrid genotype in six diverse environments by serial passaging and then used genome resequencing of evolved clones to determine evolutionary mechanisms of adaptation. The evolved clones generally increased fitness after passaging, accompanied by an average of 3.3 point mutations, 2.9 loss of heterozygosity (LOH) events, and 0.7 trisomic chromosomes per clone. LOH occurred through nondisjunction of chromosomes, crossing over consistent with break-induced replication, and gene conversion, in that order of prevalence. The breakpoints of these recombination events were significantly associated with regions of the genome with lower sequence divergence between the parents and clustered in subtelomeric regions, notably in regions that had undergone introgression between the two parental species. Parallel evolution was observed, particularly through repeated homozygosity via nondisjunction, yet there was little evidence of environment-specific parallel change for either LOH, aneuploidy, or mutations. These data show that AD hybrids have both a remarkable genomic plasticity and yet are challenged in the ability to recombine through sequence divergence and chromosomal rearrangements, a scenario likely limiting the precision of adaptive evolution to novel environments.

## Introduction

Hybridization reunites genomes that diverged via speciation. Following speciation, genomes from divergent populations have fixed nucleotide differences as well as usually larger-scale genomic changes such as chromosome rearrangements. Typically, hybridization generates individuals of low fitness due to genetic incompatibilities in the merging genomes, such as Dobzhansky-Muller genic interactions displaying negative epistasis (Orr and Turelli 2001; Mallet 2007). If structural differences exist between the chromosomes of the hybridizing genomes, they may impair proper meiosis and contribute to full or partial sterility. Overall, there exists a general relationship between genetic divergence, genome rearrangements, hybrid fitness, and reproductive isolation that is still poorly understood (Comeault and Matute 2018).

On the other hand, hybridization has the potential to facilitate adaptive evolution by bringing together novel allele combinations across loci that may produce novel phenotypes (Rieseberg *et al*. 1999; Dittrich-Reed and Fitzpatrick 2013). Particularly in plants, hybridization has led to the formation of new species, often through mechanisms such as allopolyploidization which avoids the sterility barrier concomitant with chromosomal rearrangements (Rieseberg 2001). Hybridization in fungi is increasingly recognized as having contributed to evolutionary novelty and host-range expansion (Schardl and Craven 2003; Marcet-Houben and Gabaldón 2015; Stukenbrock 2016; Samarasinghe *et al*. 2020a). However, there are many examples of fungal hybrids that appear to be restricted to the F1 diploid stage, such as the extremophile *Hortaea warneckii* (Gostinčar *et al*. 2018), *Epichlöe* endophytes (Saikkonen *et al*. 2016), *Aspergillus latus* (Steenwyk *et al*. 2020), and emerging pathogenic *Candida* spp. (Pryszcz *et al*. 2015; Schröder *et al*. 2016; Mixão and Gabaldón 2020). Because many fungi have the ability to reproduce asexually for extended numbers of generations, the F1 hybrid genotypes/species may generate transient, successful clonal lineages that avoid segregation load and chromosomal incompatibilities that may come with meiosis (Mallet 2007). Moreover, the heterosis that occurs in diploid F1 hybrids might be particularly common, because multiple mechanisms can contribute to it, such as complementation of recessive, mildly deleterious alleles, overdominance, and epistasis (Shapira *et al*. 2014).

Despite the potential for overall heterosis in F1 hybrids, it is likely that such hybrid genotypes contain a number of alleles or allelic interactions that reduce fitness, such as incompatibilities driven by negative epistasis, underdominance, or gene copy number imbalance. Given this situation, it is also likely that hybrids could undergo minor adaptive changes in genotype through mitotic processes such as chromosome copy number variation and loss of heterozygosity (LOH), which would allow genotypic change without wholesale meiotic shuffling (Bennett *et al*. 2014; Samarasinghe *et al*. 2020a). LOH occurs through a number of mechanisms, including gene conversion, nondisjunction, deletions, or crossing over (St Charles *et al*. 2012; Symington *et al*. 2014). Overall, while it is clear that asexual F1 hybrids are commonly observed in fungi, it is unclear how often and by what means they utilize their allelic diversity for adaptation and relief of incompatibilities and whether these mechanisms can allow asexual hybrids to persist for a long time despite Muller’s Ratchet (Gabriel *et al*. 1993).

*Cryptococcus* is a model basidiomycete yeast that causes opportunistic infections particularly in immunocompromised hosts. Most infections result from colonization by *C. neoformans* (serotype A) with infection by *C. deneoformans* (serotype D) rare (Cogliati 2013). These two species are diverged by at least ∼24.5 mya (Xu *et al*. 2000b). In both clinical and environmental settings, a large percentage (∼8%) of *Cryptococcus* isolates are hybrid strains, particularly those formed between *C. neoformans* and *C. deneoformans*, often referred to as AD hybrids (Samarasinghe and Xu 2018). Studies with AD hybrids show variable virulence and hybrid vigor relative to parental species, however, evidence for transgressive segregation in pathogenicity relevant traits such as resistance to UV irradiation, growth at 37 C, and resistance to antifungal drugs has been observed (Litvintseva *et al*. 2007; Lin *et al*. 2008; Vogan *et al*. 2016; Samarasinghe *et al*. 2020a).

Genomic plasticity is apparent in AD hybrids, but it is unclear which mechanisms are most important for adaptation. Most AD hybrids are close to diploid and, although they produce sexual spores in the lab, these appear to undergo an aberrant form of meiosis (Lengeler *et al*. 2001; Sun and Xu 2007; Hu *et al*. 2008; Vogan *et al*. 2013). The two species are 85-90% divergent at the sequence level, and they are known to have karyotype differences (Kavanaugh *et al*. 2006; Sun and Xu 2009). The dependence on sequence similarity for homology-based recombination, such as the classical double strand break repair pathway (DSBR) is likely to be inhibitory to mitotic and meiotic recombination at this level of sequence divergence (Datta *et al*. 1997; Opperman *et al*. 2004; Hum and Jinks-Robertson 2019). It was recently demonstrated that deletion of the DNA mismatch repair gene *MSH2* did not increase the rate of homologous recombination in meiotic progeny of an AD hybrid cross as observed in most other species but did increase viability through increased aneuploidy, a result consistent with either sequence dissimilarity or genetic incompatibility limiting recombination in *Cryptococcus* AD hybrids (Priest *et al*. 2021). Reduced recombination may explain why genomic scale studies of naturally occurring AD hybrids typically have primarily detected LOH of entire chromosomes (Hu *et al*. 2008; Li *et al*. 2012; Rhodes *et al*. 2017). A recent experimental evolution study of an AD hybrid genotype showed that LOH of Chr. 1 could be rapidly stimulated if hybrids were grown in the presence of the antifungal fluconazole, which has a major resistance allele encoded on Chr. 1 of the *C. neoformans* parent (Stone *et al*. 2019; Dong *et al*. 2020). In this same study, LOH was shown to occur at a rate of 6 x 10^−5^ per locus per mitotic division, and the distribution of LOH was unequal across chromosomes. Little is known about the extent to which more spatially restricted forms of LOH, such as gene conversion and mitotic crossing over have contributed to LOH in natural AD hybrids. The precise mechanisms of LOH in most studies are masked due to low numbers of markers utilized for genotyping, making it unclear whether the high level of sequence diversity inhibits homologous recombination in *Cryptococcus* AD hybrids through anti-recombination mechanisms.

In this study, we experimentally evolved AD hybrids to six distinct environments to test which mechanisms of genomic change are involved in rapid adaptation. Using replicate populations, both in liquid media and a model host (larvae of the waxworm moth *Galleria mellonella*), we tested for evidence of parallel mutation, LOH, or copy number variation across genes and chromosomes. Because of the divergence between parental alleles, we hypothesized that unlike in *Candida albicans* (Ene *et al*. 2018) and *Saccharomyces cerevisiae* (James *et al*. 2019) LOH would primarily be through nondisjunction of entire chromosomes rather than via mechanisms that involve homologous recombination (e.g., gene conversion or crossing over). Finally, we designed the experiment so that it would test whether particular environments would favor one parental genotype than the other as previous investigations have hinted at (Hu *et al*. 2008; Samarasinghe *et al*. 2020b). We found that whole chromosome LOH and aneuploidy were common among the evolved clones in addition to de novo point mutations. LOH events and aneuploidy demonstrated parallel evolution across independent cultures, however, these parallel events were mostly environment agnostic. In concert with the demonstration that the evolved strains have increased fitness, these data are consistent with LOH occurring to reduce genetic incompatibilities between the two parental species (Vogan and Xu 2014).

## Materials and Methods

### Strains

We initiated evolution starting with a single diploid hybrid strain generated by crossing *C. neoformans* haploid strain YSB121 (KN99 *MAT***a** *aca1::NEO ura5 ACA1-URA5*) with *C. deneoformans* strain SS-C890 (*MATα NAT* ectopically inserted into a JEC21 genome and later determined to be inserted in the sub-telomeric region of the left arm of Chr. 1). Over 100 diploid products resistant to NAT and G418 (conferred by *NAT* and *NEO* genes present in the two haploid parents) were identified that also produced extensive filamentation on V8 agar, a phenotype characteristic of cells that are heterozygous that the mating type locus and have both **a** and *α MAT* alleles. These were genotyped at 11 PCR markers across 6 chromosomes plus the *MAT* locus (**Suppl. Table 1**) to identify highly heterozygous diploid offspring. The genotypes of these markers were determined by size differences or presence/absence of a band (*STE20* locus) using agarose gel electrophoresis. One of the diploids (SSD-719) heterozygous at all markers was chosen as the ancestral genotype. A near isogenic strain (SSE-761) was generated to serve as a reference competitor strain by random integration of a GFP-Nop1-*HYG* cassette into SSD-719 (Park *et al*. 2016).

### Passaging in vitro

Experimental passaging in five media types was conducted in parallel with a *Saccharomyces cerevisiae* evolution experiment (James *et al*. 2019). Briefly, 6 replicate populations of 500 μl were inoculated with the hybrid ancestor strain, grown at 30 °C with shaking, and transferred daily with 1/50 dilution into new media. The media types were yeast peptone dextrose (YPD), beer wort, wine must, YPD + high salt (1 M NaCl), and minimal medium plus canavanine (James *et al*. 2019). Every fifth passage, 25 μl of each population was subsampled for archiving in 25% glycerol at -80 °C. At the end of 100 days, one clone was isolated from each population by streak plating, and the remaining population archived.

### Passaging in vivo

We utilized last instar larvae of *Galleria mellonella* for the infection model with inoculation and incubation following (Phadke *et al*. 2018). Six replicate populations of 5 larvae were founded, with each larva injected with 10^7^ cells. Larvae were incubated 5 days with ambient temperature and lighting. The larvae were then crushed in 50 ml sterile water and 50 μl were plated onto three 15 cm plates of YPD containing 200 mg/L G418 and 200 mg/L ampicillin. After 72 hr of incubation, the plates containing on the order of 10^3^-10^4^ small colonies were scraped and pooled into 50 ml of sterile water, the cell density was adjusted to the appropriate level (10^6^/μl) for inoculation, and the next population of 5 larvae was inoculated. After 10 passages in this manner, a single clone from each population was isolated for analysis.

### Fitness estimation

We estimated fitness of our evolved strains using both growth curves and competition with a reference strain. Growth curves were estimated using a Bioscreen C (Growth Curves USA) following James *et al*. (2019). Competition-based fitness was measured by comparing proportions of unlabeled cells (evolved) with a near isogenic competitor strain (SSE-761) engineered to express GFP. Mixtures of the competitor strain and evolved strain were analyzed for green fluorescent events to non-fluorescent events both before and after growth on a iQue Screener PLUS (Intellicyt) according to James *et al*. (2019). Each pairwise competition was run using 8 replicate populations.

Fitness was also estimated in the *Galleria*-passaged clones as well as two clones isolated from beer wort. *In vivo* fitness of the evolved strains was estimated by co-injection of evolved with competitor strain SSE-761 as a combination of 2 x 10^6^ cells in 10 μl into each of 5 larvae. After incubation for 48 hrs at ambient temperature, larvae were crushed in sterile water, filtered through a 40 μm filter, and plated on two media types: YPD media containing G418 (200 mg/L), ampicillin (200 mg/L), and chloramphenicol (25 mg/L), and YPD media containing the same antibiotics plus hygromycin (200 mg/L) and incubated at 30 °C until colony development. The latter media type only permits growth of the competitor strain while the former allows growth of both evolved and competitor. We also plated the mixture of evolved and competitor strains on the two media types to measure the ratio of the genotypes pre-injection. All experiments were repeated three independent times. Colony sizes varied considerably, and in order to count the colonies in a rapid, objective manner, we used the program OpenCFU (Geissmann 2013) using a threshold of 5 and a minimum radius of 20. By comparison of the ratios of the two genotypes pre- and post-infection, fitness can be measured. Fitness (*F*) values were calculated using colony counts on G418-only media pre- (*G_pre_*) and post-infection (*G_post_*) and colony counts on G418+hygromyin media pre- (*H_pre_*) and post-infection (*H_post_*) using the following formula:

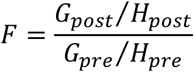

We presented the mean *F* for each clone as a relative fitness measure by dividing by the mean *F* estimate the competition between the ancestral genotype vs. congenic competitor strain.

### Genome resequencing

DNA was extracted from clones grown overnight in 2 ml of YPD following (Xu *et al*. 2000a). One clone from each population was sequenced, for a total of 35 clones (library of CB-100-4 failed to pass QC). The ancestral haploid and diploid parents were also sequenced. Because the clones evolved via passaging in *Galleria* were initiated after the *in vitro* evolution experiment using a separate subclone of the ancestor, we also sequenced the two ancestral diploid genotypes separately. Poor DNA extraction of the SSD-719 *in vivo* culture required a second DNA extraction, which resulted in additional opportunities for genomic change. Libraries for genome sequencing were constructed with dual indexing using 0.5-2.5 ng per reaction using a Nextera XT kit (Illumina). The libraries were multiplexed and sequenced on an Illumina HiSeq 2500 v4 or NextSeq sequencer using paired end 125 bp output. After analyses the expected coverage for each strain ranged from 41-147X.

### Data analyses

The full pipeline, including scripts and data sets used for analysis of the genomic data can be found at https://github.com/Michigan-Mycology/Cryptococcus-LOH-paper, with larger variant call format (vcf) files stored using figshare. Details of the variant calling pipeline using GATK 4.0.0 (McKenna *et al*. 2010) are similar to James *et al*. (2019). Briefly, data were trimmed for quality using Trimmomatic 0.36 (Bolger *et al*. 2014). The genomes of the two parental haploid strains backgrounds (JEC21 (Loftus *et al*. 2005), H99 (Janbon *et al*. 2014)) were downloaded and aligned using Harvest (Treangen *et al*. 2014). The variants from these alignments were used as a set of known variants in the base recalibration step of GATK. SNP identification for LOH and de novo mutation analyses was done using HaplotypeCaller using the H99 genome as the reference. After variant filtration for quality in GATK, SNPs were converted into a vcf file and further filtered. Genotypes were set to unknown when their depth was < 18X or > 350X, the genotype quality (GQ) was < 99, or the sites were not present as differences between the haploid ancestors observed as heterozygosity in the diploid ancestor. SnpEff 4.3K (Cingolani *et al*. 2012) was used to assign functional information to SNPs. We scanned for new mutations occurring during passaging using the criteria that they were absent in both haploid parents and ancestral diploid strain and appeared with sequencing depth > 18X and GQ >= 99. Estimating the number and length of each LOH event was done using the custom perl script calculate_LOH_lengths_crypto_100_new.pl. After identifying LOH patterns, we later verified each event by examining the read pileup and vcf file and excluded LOH events which were not supported by 100% of reads as homozygous. We removed single SNP LOH events, because they showed strong association of reads linked to only a single allele in flanking heterozygous SNPs, indicating they are likely false positives. Indels were ignored for considering LOH, which underestimates the overall amount of LOH and allelic diversity, but should not cause systematic biases in hypothesis testing.

We identified aneuploidy by mapping reads to a combined AD hybrid reference as in Priest *et al*. (2021). In this hybrid reference sequence, the *C. deneoformans* chromosomes have been reordered and reverse-complemented where necessary to better match the chromosome order of *C. neoformans*. The trimmed reads of each strain were mapped using bwa v. 0.7.15 (Li and Durbin 2009), reads of low mapping quality (-q 3) removed using samtools v. 1.5 (Li *et al*. 2009), and coverage estimated along chromosomes in 5 kb intervals with bedtools v. 2.29.2 (Quinlan and Hall 2010). The data were then plotted using ggplot2 (Wickham 2009) after removal of windows with depth of coverage > 5X the genome-wide average. Local scale copy number variation (CNV), specifically looking for deletions, was evaluated using the software smoove (https://github.com/brentp/smoove).

All statistical analyses and plots were conducted in R 3.6.2 (R_Core_Team 2019) unless otherwise specified. Most statistical tests utilized the base package; we also utilized the functions ggplot2 and gridextra for graphical output (Auguié 2017).

### Data availability

Strains and plasmids are available upon request. The full pipeline, including scripts and data sets used for analysis of the genomic data, additional data files, and code for reconstructing figures can be found on Github (https://github.com/Michigan-Mycology/Cryptococcus-LOH-paper). Raw sequence data has been deposited into NCBI’s SRA archive under BioProject: PRJNA700784, with the accession numbers SRX10055646-SRX10055684.

## Results

### Fitness estimates suggest adaptation during passaging

We evolved a single AD hybrid genotype (SSD-719) in 5 *in vitro* environments by serial transfer for 100 days and also evolved the genotype through 10 serial passages in waxworm larvae. The experiment was run in parallel with an investigation of microevolution in *Saccharomyces cerevisiae*, and while the media types are more typical for brewing yeast, they served to represent broadly diverse environments that would require adaptation by the laboratory-created *Cryptococcus* hybrids. Each environment was represented by 6 replicate populations, and from each of these evolved populations we isolated individual clones, determined whether they had evolved a higher fitness relative to the ancestral genotype, and sequenced their genomes. The 6 clones isolated after evolution in each *in vitro* environment generally showed increased fitness relative to the ancestral genotype (**Figure 1**). Maximal population density (EOG) and competitive fitness mostly increased for all strains and environments, whereas doubling time was lower for clones evolved in high salt or canavanine environments.

**Figure 1.**
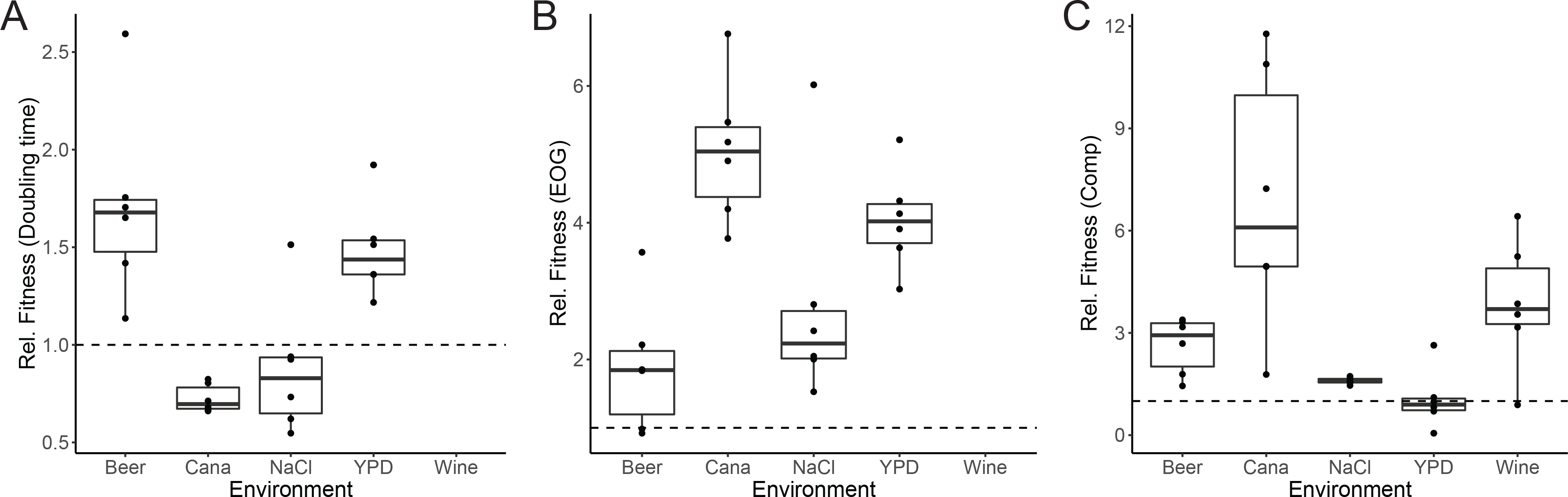
Clones evolved in *in vitro* conditions mostly showed fitness gains regardless of means of estimation. Shown are values of relative fitness comparing a clone grown in the same media in which evolution occurred relative to the fitness of the ancestral genotype SSD-719 (dashed line). A) Fitness measured using doubling time. B) Fitness measured using EOG = Efficiency of Growth (i.e., carrying capacity). C) Fitness measured using relative competitive growth in the presence of a marked ancestral strain. Each dot represents an individual clone. Boxes indicate the 25-75% percentiles, bars indicate medians, and the whiskers extend to 1.5 x the interquartile range. Cana=minimal media with a sublethal concentration of canavanine (2 mg/L) plus arginine (4 mg/L), and NaCl=complete media with 1.0 M NaCl. Wine strains were not tested by growth curve analysis due to opacity of the medium which prevented accurate readings by spectrophotometry.

The virulence and relative fitness of the *G. mellonella* passaged strains was estimated for the 6 clones that had been serially passaged 10 times in the same host compared with the ancestral diploid genotype and isolates that had been evolved in beer wort. The larval passaged strains showed enhanced survivorship (one-way ANOVA, [F(8,18)=3.59, P=0.0115]) compared to the beer wort evolved and ancestral strain as measured by genotype recovery after co-injection with a marked strain (**Figure 2**).

**Figure 2.**
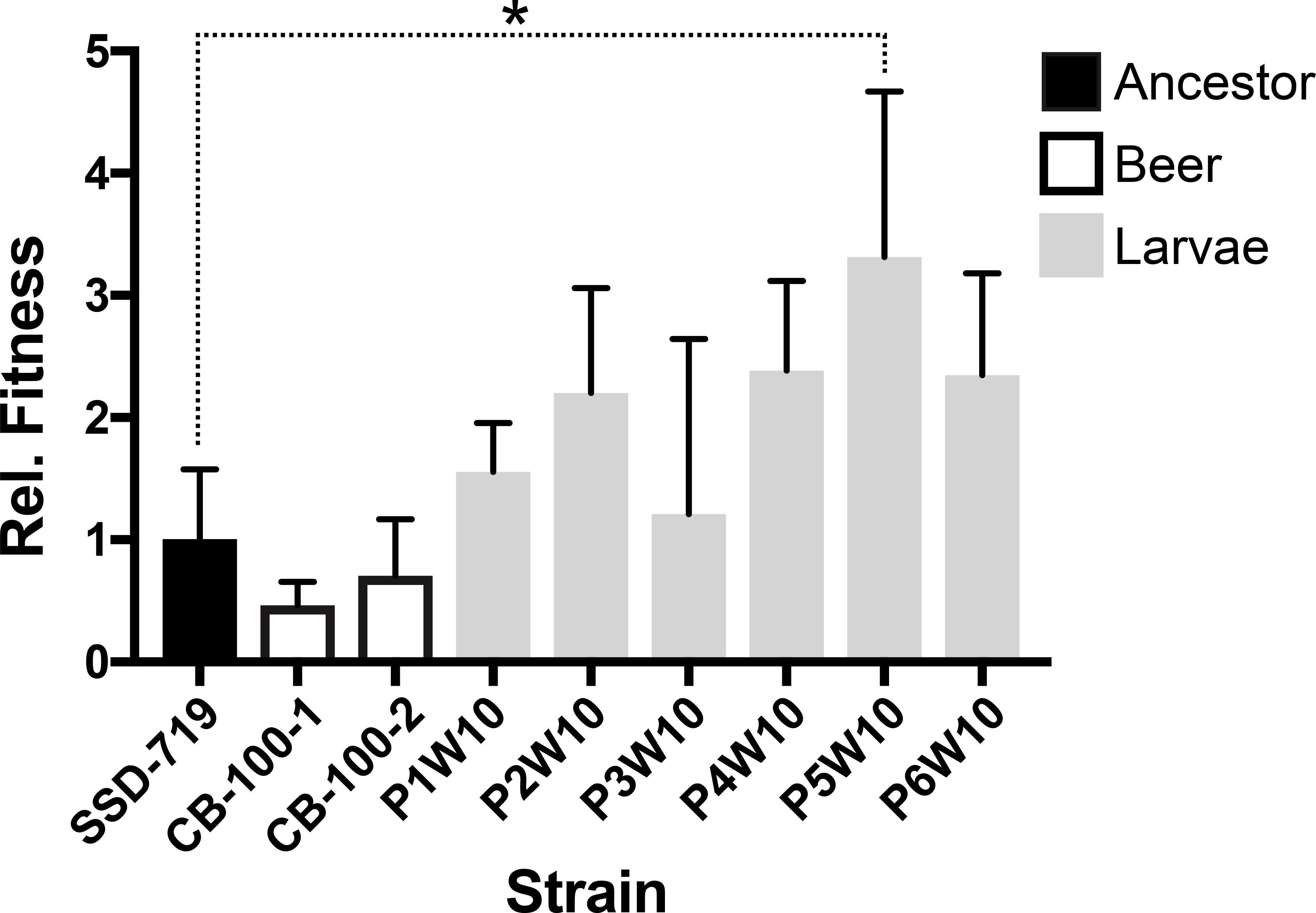
Strains passaged in larvae showed overall greater fitness than the ancestor as measured by recovery after 2 days growth post-coinjection with a common competitor strain (SSE-761). Three replicate experiments were conducted for each strain, and values indicate mean (+1 sd). All values are shown relative to the fitness of SSD-719 (ancestor) which was set to 1.0. Post-Hoc comparisons of the fitness between each evolved strain and the ancestor by Fisher’s Least Significant Difference test found only the difference between SSD-719 and P5W10 to be significant (P=0.0039). P4W10 and P6W10 were significantly different from SSD-719 to a lesser degree (P=0.06 and P=0.07, respectively).

### Parallel LOH was observed mostly as whole chromosome loss in a non-environment specific manner

LOH was readily apparent at the chromosome scale (**Figure 3**), suggesting that most strains had undergone chromosome-scale LOH events as well as multiple smaller events. We estimated that strains had undergone an average of 2.9 LOH events after 100 days, encompassing an average of 2.26 Mbp. There was a significant difference between the average number of LOH events across environments (Kruskal-Wallis test, X^2^(5) = 18.26, *p* = 0.026), but these differences were driven mostly by low LOH numbers in larval passaged strains (beer wort: 2.8, canavanine: 4.2, high salt: 1.8, wine must: 2.7, YPD: 4.7, larvae: 0.8). Most of the LOH events were the result of whole chromosome events, with the second most common being long distance LOH consistent with break-induced replication (BIR) observed as crossing over not associated with gene conversion, followed by gene conversion, and then deletions (**Figure 4**). Crossover and deletion events averaged 173 kb in total length, and gene conversions averaged 7.4 kb.

**Figure 3.**
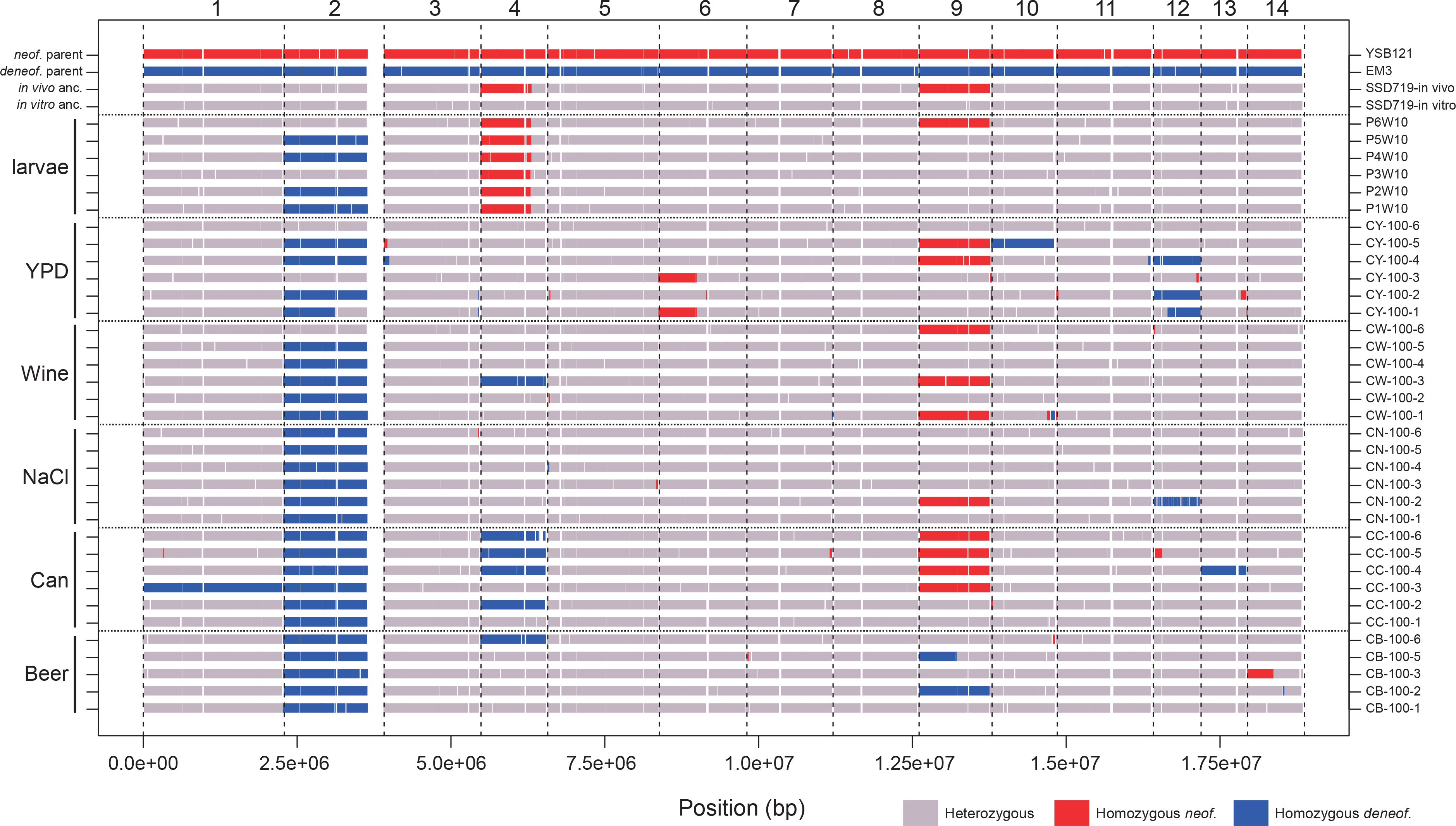
Landscape of LOH across the 14 chromosomes of the evolved clones. Shown are genotypes at 10 kb resolution of the strains, with media type shown on the left margin and strain name on the right margin. The ancestral genotype of the *in vivo* (larval) passaging was different from the *in vitro* (other media types), because the experiments began at different times. The reference genome coordinates are from the *C. neoformans* parent. White regions represent unmapped, typically repetitive regions of the genome, or regions of the genome that had undergone LOH in the ancestral diploid strains. In particular, ∼260 kb of the right arm of Chr. 2 underwent an LOH event in the ancestor before passaging.

**Figure 4.**
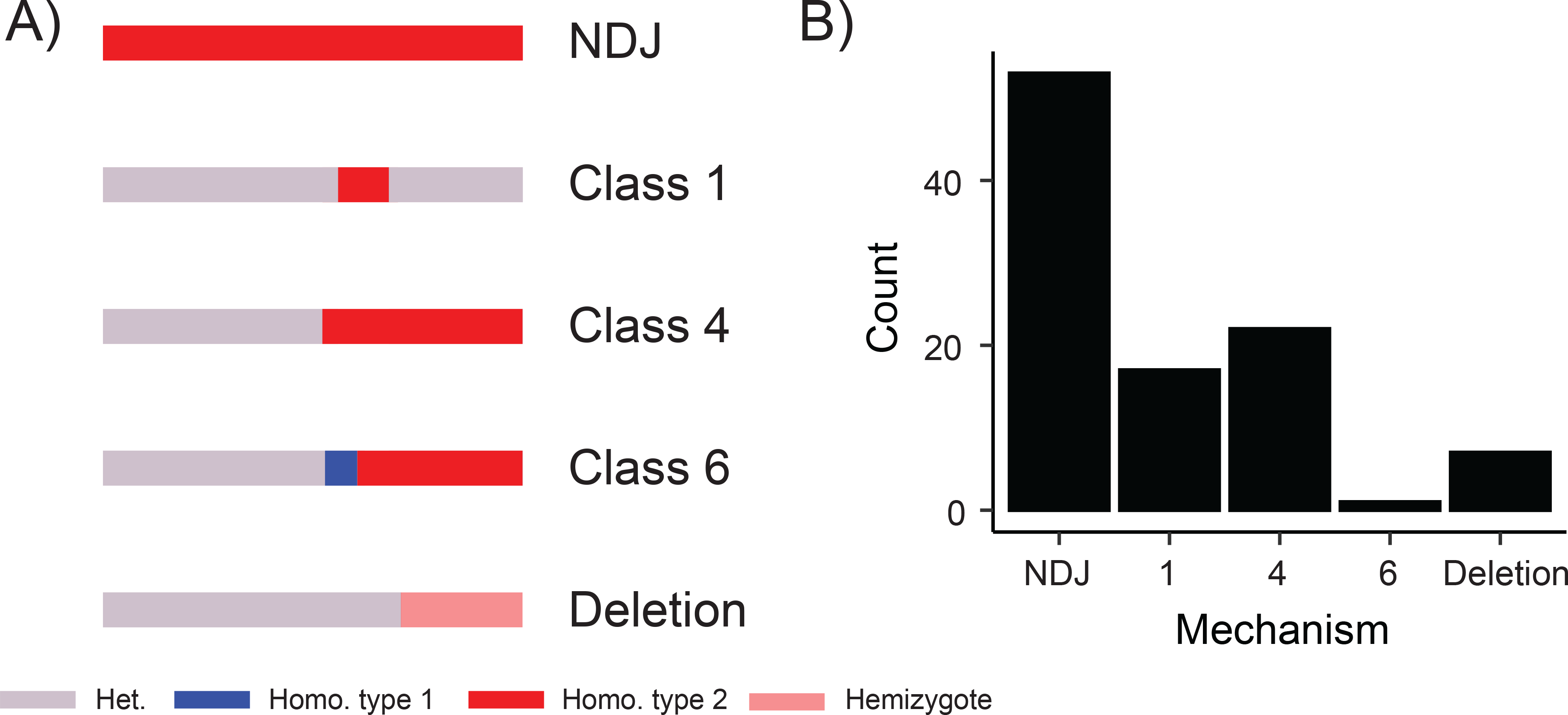
Summary of detected LOH mechanisms across the 35 evolved hybrid clones. (A) A depiction of proposed LOH mechanisms according to schema defined by St. Charles (St Charles *et al*. 2012), and (B) Tallied numbers of events observed among the hybrid clones.

We looked for evidence that specific LOH events showed parallel evolution across independently evolved clones. We specifically noted parallel changes as regions of the genome undergoing LOH in 3 or more clones. Among the LOH events, 11 were found in more than 3 clones, including whole chromosome or long distance LOH on Chr. 2, 4, 9 and 12 (**Figure 3**). The other regions were limited to only 3 clones. There was no evidence for any of the LOH events being more predominant in particular environments.

We asked whether whole-chromosome LOH events favored one parental type over another, which was obvious for Chr. 2 in which LOH exclusively led to retention of the *C. deneoformans* homolog. For the 4 chromosomes (2, 4, 9 and 12) with 4 or more independent LOH events we tested whether the allelic distribution of homozygotes was non-random or skewed towards one parent. Although events at most chromosomes favored one parental genotype over another, only the Chr. 2 pattern was significant (P=8.2e-06; Fisher’s exact test).

We then tested whether one of the parental genotypes was globally favored over the other across isolates by asking whether the ratio between SNP loci that are homozygous for the *neoformans* and *deneoformans* alleles, respectively, deviated significantly from a random 1:1 null hypothesis. Overall, we found that the mean frequency of *neoformans* alleles in LOH regions across all evolved clones was 0.234 (CI: 0.115-0.353), which was significantly different from the null hypothesis (t(34) = -4.61, p = 7.0e-5). However, if Chr. 2 was excluded, the mean frequency of *neoformans* alleles in LOH regions was 0.639 (CI: 0.488-0.791), which was not significantly different from the null expectation (*t*(34) = 1.88, *p* = 0.070). These data suggest there is no signal of a favored parental allele arising from the LOH process with the exception of LOH favoring retention of the *C. deneoformans* Chr. 2 homolog.

Using the rate of LOH events uncovered and their sizes, we estimated a rate of 0.005 events/generation, 4.0 kb/generation, and 2 x 10^−4^ per bp per generation. Dong et al. (2019) estimated a rate of LOH of 6 x 10^−5^ per locus per mitotic division with 33 loci in *Cryptococcus* AD hybrids, leading to an estimation of 0.002 LOH events per generation.

### Prevalence of aneuploidy across passaged isolates

Aneuploidy leading to chromosomal copy number variation is a rapid means with which pathogenic fungi generate adaptive genetic variation (Bennett *et al*. 2014; Beekman and Ene 2020). We tested for the presence of both whole chromosomal and partial chromosomal aneuploidy by using the density of reads mapped to a hybrid version of the genome, which we found to give a clearer picture of depth of coverage than solely using one species reference genome. The mapping data also provided strong confirmation of our LOH results. Most of the strains had mapping patterns that fell into distinct chromosome counts of 0-3, with a few cases in which average mapping density appeared to be intermediate, a result consistent with ongoing chromosome loss in strains used for genome sequencing (**Figure 5**). We found that the evolved strains often showed aneuploidy on multiple chromosomes (**Figure 6**; **Suppl. Figures 1-6**). In all cases, aneuploidies were trisomies and no evidence of monosomies was found. Particular chromosomes (Chr. 2, 4, 9, 10, and 11) were likely to display aneuploidy and were strikingly overlapping with the same chromosomes that had also undergone whole chromosome LOH (**Figure 3**). Among the evolved lines, there was a 2:1 bias towards trisomies that involved extra copies of the *C. deneoformans* homeologue. Some aneuploidies were also present in the ancestral diploid but not haploid genotypes (**Suppl. Figure 6**). For example, Chr. 9 was trisomic in all larval passaged clones, but was also observed in the *in vivo* diploid ancestor, although most passaged strains maintained heterozygosity on Chr. 9 unlike the ancestor. Chr. 3 of *C. deneoformans* was duplicated in the *in vitro* ancestor, but most of the evolved strains did not have this aneuploidy (**Figure 6**). Clone CW-100-4 evolved in wine must revealed a complex pattern of chromosome ploidy consistent with either multiple subpopulations within the strain with different aneuploidies or possibly a primarily tetraploid strain (**Suppl. Figure 3**). Overall, there were 25 aneuploidies that uniquely appeared after passaging, for 0.7 events per clone.

**Figure 5.**
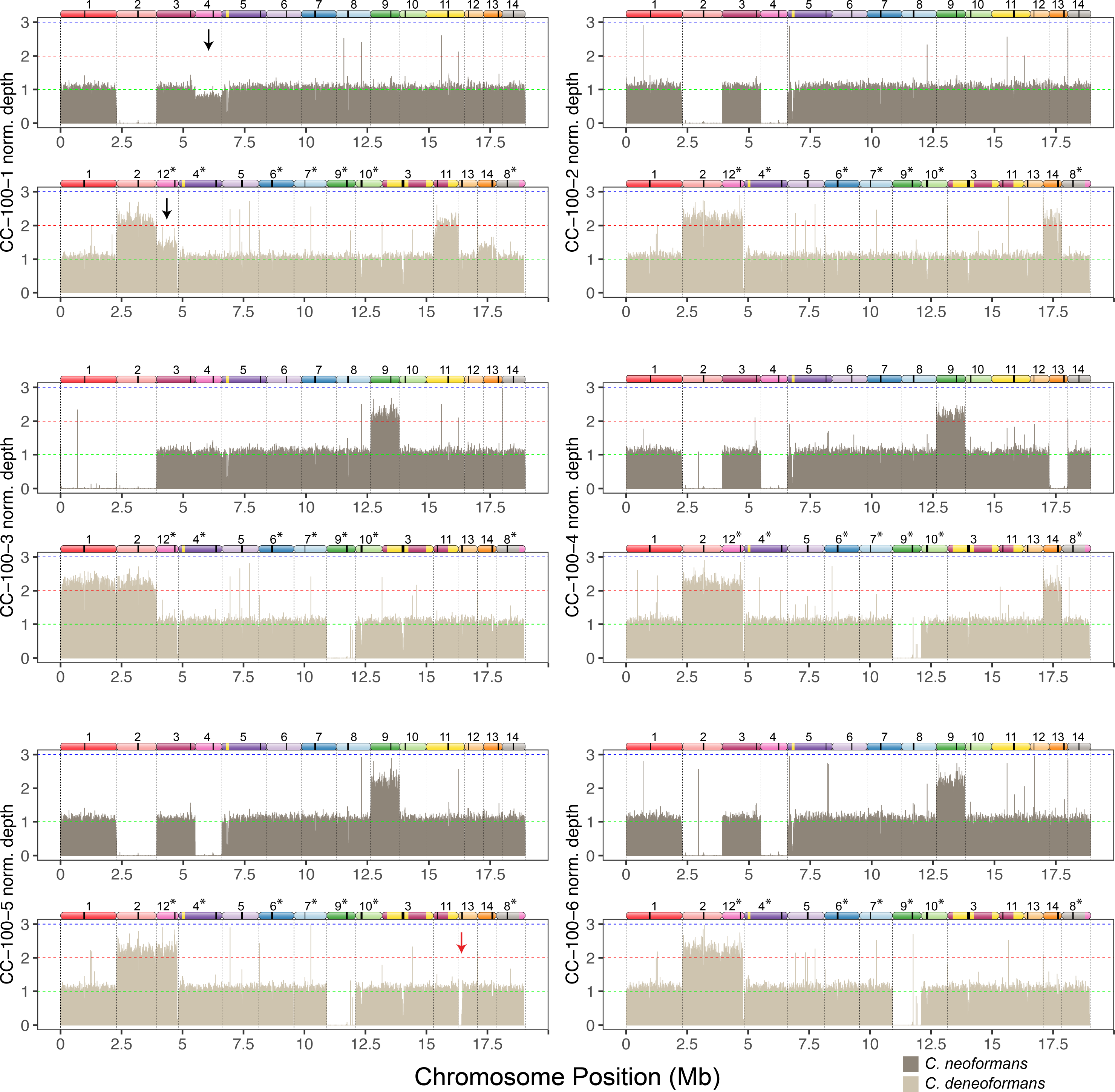
Mapping of reads to a hybrid reference genome of strains evolved in canavanine reveals trisomies and whole chromosome LOH. Bars show mean depth in 5 kb intervals that have been normalized by dividing the depth of each interval by the mean coverage for each strain. For each strain the top panel shows mapping to the *C. neoformans* reference genome, and the bottom panel shows mapping to the *C. deneoformans* reference genome. Diagrams of homeologous chromosomes are shown above each reference genome where colors reflect homology and black stripes indicate centromeres. *C. deneoformans* chromosomes were rearranged to better reflect homology, and asterisks indicate chromosomes that have been reverse-complemented. Black arrows in strain CC-100-1 indicate loss of the *C. neoformans* Chr. 4 copy and gain of the homeologous *C. deneoformans* Chr. 12 copy, likely indicating LOH ongoing during the growth of the strain for DNA extraction. Red arrow in CC-100-5 indicates a region of the left arm of *C. deneoformans* Chr. 13 that had undergone apparent deletion.

**Figure 6.**
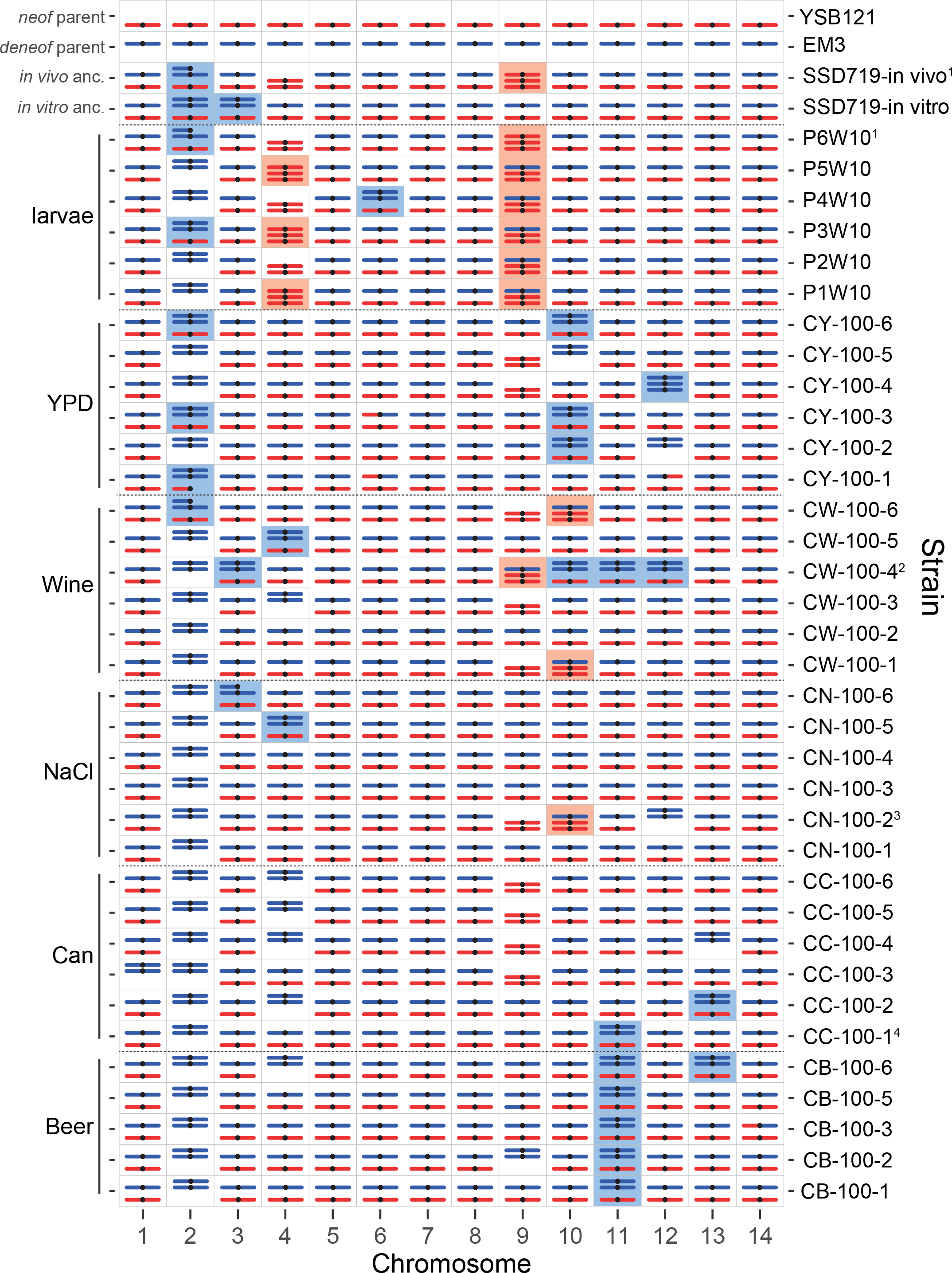
Chromosome composition of the 35 evolved clones and the ancestors reveals at least one trisomy in most strains. Partial aneuploidies are indicated as partial chromosomes that are not shown to scale. Shaded boxes indicate partial or complete aneuploidy for *C. neoformans* (red) or *C. deneoformans* (blue) homeologues. Recombined chromosomes are not indicated with the exception of a few chromosomes with larger exchanges. Footnotes indicate strains which had evidence for mixed genotype composition; all notes refer to *C. neoformans* chromosome numbers. 1. SSD719-in vivo and P6W10 showed a pattern consistent with a mixed population of trisomies for *C. neoformans* or *C. deneoformans* Chr. 6. 2. Strain CW-100-4 has a pattern consistent with either an equal mixture of two strains with different aneuploidies or a strain that is largely tetraploid. 3. Strain CN-100-2 has a pattern consistent with *C. neoformans* Chr. 12 apparently present in small subpopulation. 4. CC-100-1 has a pattern of apparent active loss of *C. neoformans* Chr. 4 and duplication of the corresponding homeologue *C. deneoformans* Chr. 12 (arrow in Figure 5). For additional details on the mapping see depth of coverage plots (Figure 5; Suppl. Figures 1-6).

Chr. 2 shows a complex history after the creation of the hybrid ancestor by mating in the lab. Both *in vitro* and *in vivo* ancestors show a deletion on the right arm of *C. neoformans* Chr. 2, spanning ∼260 kb. This was accompanied by a duplication of the entire *C. deneoformans* homeologue in the *in vitro* ancestor and a partial duplication of the *C. deneoformans* homeologue in the *in vivo* ancestor. Most of the *in vitr*o evolved strains ultimately lost the *neoformans* Chr. 2 homeologue, however, it was retained in clones CW-100-6, CY-100-3, CY-100-6, and in part in CY-100-1 (**Suppl. Figures 3-4**). The loss of the *C. neoformans* Chr. 2 homeologue may have been advantageous to the strains; strains CY-100-3 and CY-100-6 that retained the *C. neoformans* homeologue had the two lowest fitness values for YPD evolved strains in two fitness measures (doubling time and relative fitness versus competitor). Most of the trisomies did not show an environment specific pattern, however, trisomy derived via duplication of the Chr. 11 *C. deneoformans* homeologue was present in all beer wort evolved strains and one strain evolved in canavanine. This trisomy was significantly associated with beer wort (Fisher’s exact test, P=8.6e-05), suggesting this association may have been driven by an environment-specific fitness advantage of this extra chromosome.

Chromosome plots and a structural variant detection method (smoove) were also employed to detect partial aneuploidy occurring via partial chromosomal deletion. Seven LOH events were consistent with deletions (2 identified by smoove and 5 visually from relationship of coverage with LOH events). Overall, these data suggest that most strains are not diploid but instead aneuploid with duplication of the *C. neoformans* homeologue more frequently involved in trisomies of particular chromosomes.

### Absence of parallel evolution by point mutations

We found 117 high quality *de novo* mutations at 93 SNP loci specifically occurring in the evolved clones that were absent from both haploid and diploid ancestors. The mean number of mutations per strain was 3.3 (range 0-10), and mutations were found on every chromosome and in every environment type. The majority of the mutations were inside exons (81.7%), with the remaining intronic (10.8%), or intergenic (8.5%). Most of the mutations in exons had consequences on the protein sequence, with 64.5% missense, 6.6% nonsense, and 28.9% synonymous (**Suppl. Table 2**). Of 117 mutations only 4 of them were found as homozygous genotypes. Because a number of chromosomes have undergone whole chromosome or long distance LOH, the fact that most mutations are heterozygous implies that whole chromosome LOH likely occurred early on during the evolution of the populations, followed by mutations. Of the 93 *de novo* single nucleotide mutation loci, only 3 were found in 3 or more strains. Three loci showed 3, 7, and 8 strains bearing the mutation. These were all mutations of low impact and may represent variation in the initial founding population that became fixed during evolution. Mutations with protein-altering effects were found in diverse genes, including transcription factors, plasma-membrane proton-efflux P-type ATPase, and many others. A test for enrichment of Gene Ontology terms among the protein-altering mutations using clusterProfiler (Yu *et al*. 2012) identified lipid metabolic process as the most enriched term, although this enrichment is not statistically significant after correction for multiple testing (FDR corrected P = 0.103).

### Relationship of LOH break points to sequence divergence and mutation

The ability to undergo crossing over is known to be influenced by anti-recombination mechanisms that prohibit recombination due to sequence divergence. Such a mechanism would favor mitotic recombination breakpoints in regions of lower divergence. We found no relationship between heterozygosity of particular chromosomes and the number of crossover events observed (r(12) = 0.083, p = 0.78; **Suppl. Table 3**). By plotting the locations of mitotic recombination breakpoints and mutations relative to chromosomal divergence (**Figure 7**), however, we identified a clear pattern that many of the breakpoints are sub-telomeric, and sub-telomeric regions often have lower divergence between the two parents. We tested whether the heterozygosity near crossing over breakpoints was depressed relative to random breakpoints produced in simulated data sets. In total there were 47 independent breakpoints observed (only the left breakpoint was used for gene conversion events). For windows of size 1000 bp centered around the breakpoints, the mean post-filtering heterozygosity of the windows was 0.026. From 1000 simulated data sets, the mean simulated divergence near random breakpoints was much higher (0.045), with a range of 0.037-0.051, providing a highly significant (P<0.001) difference between heterozygosity near observed breakpoints to randomly chosen ones. Varying the window size from 30-10^5^ bp revealed that heterozygosity near LOH breakpoints was lower than the random expectation for every case (P<0.001). This result suggests that recombination in *Cryptococcus* AD hybrids is favored in broad regions of lower divergence between recombining genomes.

**Figure 7.**
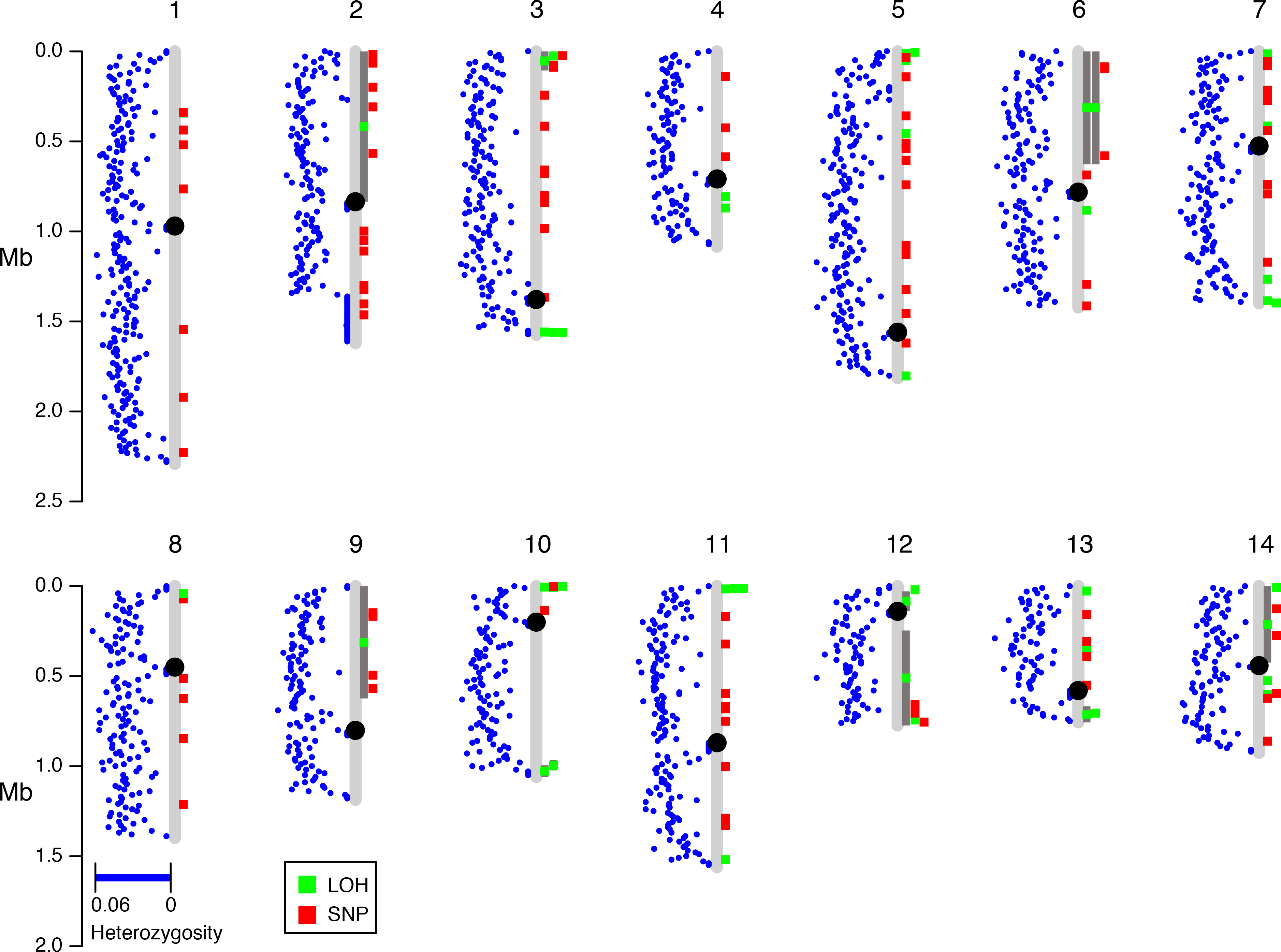
Locations of mitotic recombination breakpoints are non-random across the genome. Blue dots indicate the average heterozygosity in 10 kb windows between *C. neoformans* and *C. deneoformans* parental genomes that passed our filtering strategies. Green boxes indicate locations of LOH events, and when accompanied by dark grey bars they indicate longer LOH tracts.

We investigated the breakpoints further to understand if there were sequence characteristics that promoted recombination and identified hotspots at two regions of the *Cryptococcus* genome where ancestral introgression likely occurred between the two parental lineages (Kavanaugh *et al*. 2006). The smaller of the introgression events encompasses ∼12 kb of the left arm of *C. neoformans* Chr. 10 which is found on the right arm of Chr. 13 in *C. deneoformans*. This small region of ∼5% sequence divergence harbored 3 of the 47 independent breakpoints. The second, larger introgression event is a 40 kb region found in a sub-telomeric region of the left arm of *C. neoformans* Chr. 5 which was apparently the parent of a non-reciprocal introgression and translocation into *C. deneoformans*. Strikingly, 4/6 non-whole chromosome LOH events occurring on *C. neoformans* Chr. 5 have a crossover event in this “identity island”. Two of these events are gene conversions with one end near the boundary of the island where one partial copy of the non-LTR retrotransposon Cnl1 is present.

In *Candida albicans*, a positive relationship has been observed between mitotic recombination and *de novo* mutation (Ene *et al*. 2018), indicative of recombination-induced mutagenesis as observed in meiosis in yeast and other organisms (Arbel-Eden and Simchen 2019). We tested for such a relationship by comparing the relative distance of observed mutations to crossover locations to a simulated set of random mutations in the genome (1,000 simulations). This result supports the overview in **Figure 7** that suggests there is no clear relationship between *de novo* mutation and crossover events in *Cryptococcus* AD hybrids (P = 0.508).

## Discussion

Mutations are the primary way in which clonal organisms evolve. However, there are additional avenues for genetic change in clonal organisms referred to as genome plasticity, which are particularly available to polyploid hybrid genomes (Albertin and Marullo 2012; Blanc-Mathieu *et al*. 2017; Samarasinghe *et al*. 2020a). In this study we investigated the sources of genetic variation in evolving hybrid strains of *Cryptococcus* continuously passaged in differing environments. Passaging led to adaptation as observed by increased fitness of the clones isolated at the end of the experiment relative to the ancestral genotype. Genome resequencing allowed an unprecedented look at lab-evolved *Cryptococcus* hybrids and revealed multiple mechanisms of genomic change. Genomic plasticity was observed as chromosome copy number amplification and LOH generated via nondisjunction, gene conversion, and crossing over. However, the location of these genome scale events is highly biased across the genome, occurring in particular chromosomal locations presumably deeply influenced by the history of the divergence of the two sibling species.

This experiment was conducted at the same time as a similar passaging experiment using *Saccharomyces cerevisiae* (James *et al*. 2019), and therefore the media conditions were atypical for *Cryptococcus*. Therefore, it is unsurprising that the *Cryptococcus* hybrid genotype improved significantly in fitness over the period of evolution, particularly as measured by density at carrying capacity (EOG) and in head-to-head competition with the ancestor (**Figure 1, B-C**). *Cryptococcus* AD hybrids had a lower average LOH rate (2.9 event/100 days) relative to *S. cerevisiae* (5.2 events/100 days), despite the same dilution rate on transfer and hence equivalent number of generations. Moreover, the mechanism of LOH in *Cryptococcus* AD hybrids was very different than that of *S. cerevisiae* heterozygous diploids. Specifically, LOH in *Cryptococcus* is dominated by nondisjunction of chromosomes, while LOH in *S. cerevisiae* had gene conversion as the majority mechanism and no evidence of whole chromosome LOH. Both fungi had LOH consistent with crossing over or BIR (**Figure 4**; Class 4), but *S. cerevisiae* had considerably more crossing over associated with gene conversion, consistent with the classical double strand break repair model (Symington *et al*. 2014). A major difference that may explain this pattern is that the initial heterozygosity of the ancestral diploid genotypes of *Saccharomyces* was 0.005-0.009 or one heterozygous position out of every 100. In our highly filtered *Cryptococcus* data set, we considered 5.6e05 heterozygous positions in the AD hybrids (0.03 heterozygosity), however, it is known that *C. neoformans* and *C. deneoformans* are 85-90% divergent at the sequence level (Kavanaugh *et al*. 2006; Sun and Xu 2009) making differences on the order of 10 bases for every 100.

LOH and aneuploidy was highly heterogeneous across the *Cryptococcus* genome. Whole chromosome LOH was primarily observed on chromosomes 2, 4, and 9, and trisomy was common for these chromosomes plus 10 and 11. Comparison of *C. neoformans* to *C. deneoformans* has identified 32 chromosomal rearrangements (Sun and Xu 2009). Most of these are smaller inversions or complex rearrangements, while a few of them represent large chromosomal changes, such as multiple translocations and inversions between chromosomes 3 and 11. The rearrangement between Chr. 3 and 11 had a severe and suppressive impact on long distance LOH both as whole chromosome loss or as crossover events, however these chromosomes may still play a role in adaptation through chromosomal copy number changes, and one amplification (Chr. 11 of *C. deneoformans*) was positively associated with growth in beer wort. Other larger chromosomal rearrangements likely had additional suppressive effects on the ability of the homologous chromosomes to undergo mitotic recombination. For example, no crossover events were observed on the right arm of chromosome 9, which is expected due to the large inversion between *C. neoformans* and *C. deneoformans* in this region (Sun and Xu 2009).

If sequence dissimilarity and chromosome rearrangements hinder mitotic recombination what might stimulate it? Crossover locations were highly biased towards sub-telomeric regions, and these are regions where there are fewer heterozygous mapped positions. It is worth pointing out, however, that there are multiple explanations for lower heterozygosity at the telomeric regions. One could be hemizygosity at these regions, or alternatively the enrichment of repetitive DNA elements such as transposons in these regions (Loftus *et al*. 2005), which remain unmapped with short read technology. It is clear that there is an association between transposable elements and chromosomal rearrangement and introgression (Kavanaugh *et al*. 2006; Sun and Xu 2009). We found that regions of lower divergence caused by transposable element-associated introgression were hotspots of mitotic recombination. These results allude to a scenario in which the transposable elements both speed up and slow down the speciation process by both increasing genome rearrangements but also increasing genetic similarity through introgression, which should increase the ability to recombine.

While this experiment played out over a mere ∼500 generations, naturally occurring hybrids reveal evolutionary processes occurring over much longer time periods. *Cryptococcus* AD hybrids have arisen multiple times independently and may have undergone clonal evolution for up to 2 million years (Xu *et al*. 2002; Litvintseva *et al*. 2007). The results of our microevolution study show many similarities to genomic analysis of the natural hybrids. For example, these studies have observed both whole chromosome LOH and aneuploidy (Hu *et al*. 2008; Sun and Xu 2009; Li *et al*. 2012; Rhodes *et al*. 2017). The specific chromosomes involved, however, are different. While there are shared whole chromosome LOH events between our study and others as in loss of the C*. deneoformans* Chr. 9 homolog or loss of the *MAT* encoding Chr. 4 (Sun and Xu 2009; Li *et al*. 2012; Rhodes *et al*. 2017), the general pattern across strains and studies is mostly stochastic, which contrasts with the highly deterministic pattern seen for Chr. 2 in our experimental population. A possible explanation is that the hybridization events are independent, and all of a different background than the lab generated one used in this study. Many hybrids have the VNB lineage as the *C. neoformans* parent, but in our study the *C. neoformans* parent was VNI (Litvintseva *et al*. 2007; Li *et al*. 2012). Another possibility is that the LOH events observed are both stochastic yet contingent on existing genotype. Specifically, our ancestral genotype lost the right 260 kb of the *C. neoformans* Chr. 2 before passaging. This event may have set up a scenario where further loss of the entire *C. neoformans* Chr. 2 was favored, perhaps due to genetic incompatibilities between genes on this chromosome.

Genomic plasticity via LOH and aneuploidy are predicted to be adaptive, though this must be contrasted with a more neutral model. In the case of drug resistance in AD hybrids aneuploidy and increased copy number is clearly selected for through amplification of resistance genes (Li *et al*. 2012; Rhodes *et al*. 2017; Dong *et al*. 2020). Aneuploidy may merely be a byproduct of the fact that chromosomal loss or duplication is the fastest means of achieving copy number change or homozygosity of a single gene. That selection was responsible for the rapid increase in homozygosity of Chr. 2 to the *C. deneoformans* homolog is supported by the fitness values of strains without homozygosity of the Chr. 2 *C. deneoformans* homolog. These data are also consistent with the strong bias towards the *C. deneoformans* parent of Chr. 2 in a large sample of sexually recombined progeny from an AD hybrid of similar genetic background (Sun and Xu 2007), hinting that there may exist incompatibilities between genes from the two species on this chromosome as predicted by the Dobzhansky-Muller model. Such intra- and inter-chromosomal incompatibilities have been detected in a previous study of AD hybrids, suggesting that this a plausible explanation (Vogan and Xu 2014). Alternatively, rather than epistatic interactions, dosage or RNA/protein titer could differ between products encoded by the two parental homologs, and this could provide opportunities for selection to act. For example, if there are genes not present on Chr. 2 of *C. neoformans* that exist in *C. deneoformans* or if the expression level of the *C. deneoformans* genes on Chr. 2 is more optimal, this could lead to selection favoring loss of the *C. neoformans* homolog. As Chr. 2 is the chromosome containing the ribosomal RNA (rRNA) array, we might hypothesize that either copy number or expression of these genes could be involved in rapid whole chromosome LOH.

Overall, the contrasting results in LOH rate and mechanism between *Cryptococcus* hybrids and *Saccharomyces* diploids (James *et al*. 2019; Sui *et al*. 2020) suggest that LOH in *Cryptococcus* hybrids is comprised largely of whole chromosome events in part because of the inability for *Cryptococcus* AD hybrids to undergo recombination. This evidence comes from the association between crossover breakpoints and regions of lower heterozygosity, such as identity islands shaped by introgression, an observation consistent with literature showing mitotic and meiotic recombination is reduced in species with higher heterozygosity (Opperman *et al*. 2004; Seplyarskiy *et al*. 2014; Hum and Jinks-Robertson 2019; Tattini *et al*. 2019). Hybrid linkage maps also support suppression of recombination in *Cryptococcus* AD hybrids (Sun and Xu 2007; Vogan *et al*. 2013). In contrast, intraspecific linkage maps constructed using *C. deneoformans* showed results comparable to *S. cerevisiae* (∼5 kb/cM) (Cherry *et al*. 1997; Forche *et al*. 2000; Sun *et al*. 2014; Roth *et al*. 2018), revealing that homologous recombination is not generally inhibited in the genus. Rates of mitotic recombination within *Cryptococcus* intraspecific crosses have not been measured. In our study design, we also evolved *C. deneoformans* diploids that were completely homozygous. These homozygous diploid populations showed a similar rapid increase in fitness as evolved *Cryptococcus* hybrid clones, and future genome sequencing could resolve whether most *de novo* mutations remain heterozygous as observed in hybrid clones or may have undergone LOH at a higher rate.

## Acknowledgements

This work was made possible by grants from the National Institutes of Health, National Institute of Allergy and Infectious Diseases (5R21AI105167-02, R37 AI39115-2<u>4</u>, and R01 AI50113-16). J.H. is co-director and he and T.Y.J. are fellows of CIFAR program Fungal Kingdom: Threats & Opportunities. We thank Emmi Mueller and Ellen James for technical lab support. We thank Marco Coelho for providing the hybrid reference genome and the schematics of the *Cryptococcus* chromosomes.

---

**Suppl. Table 1.**
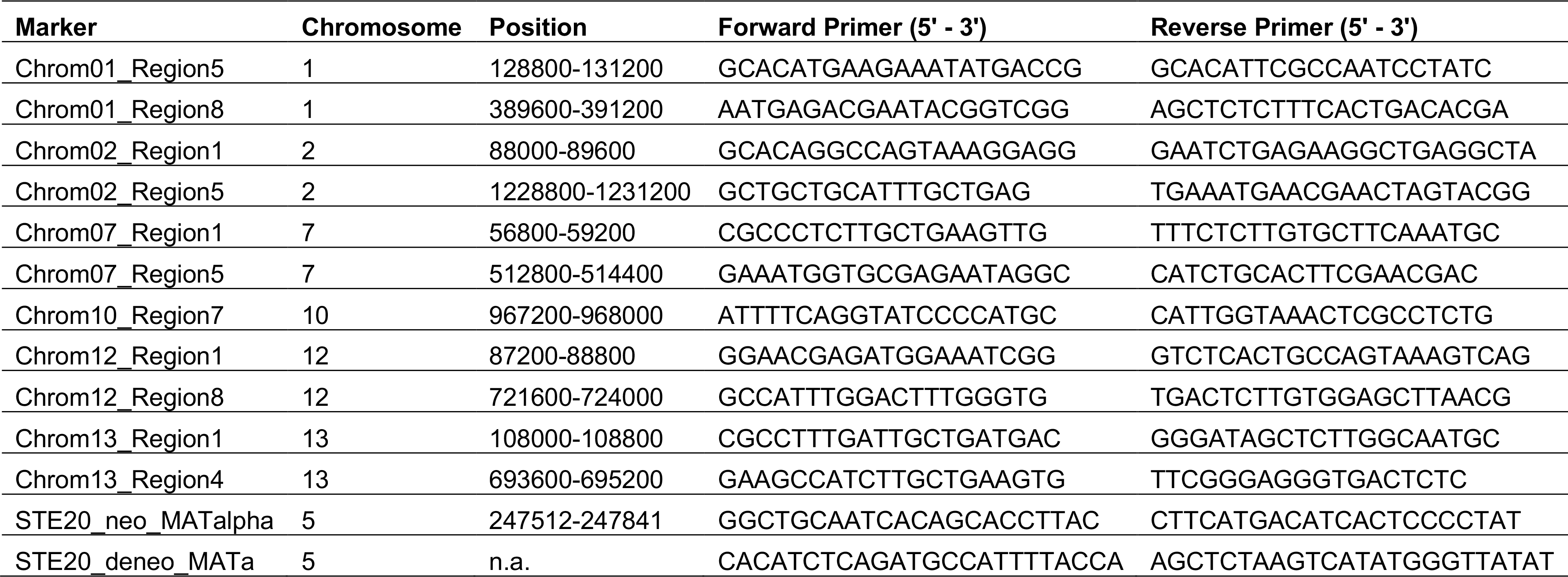
Markers used for screening mating products for heterozygosity. Chromosome numbers and positions are relative to the H99 genome.

**Suppl. Table 3.** Heterozygosity across chromosomes is not correlated with frequency of crossover events or nondisjunction (NDJ). Data are relative to *C. neoformans* H99 genome. Only one breakpoint for each gene conversion is considered.

| Chr | Size | Num SNPs | SNP rate | Num. breaks | NDJ |
| --- | --- | --- | --- | --- | --- |
| 1 | 2291499 | 72959 | 0.0318 | 1 | 1 |
| 2 | 1621675 | 39814 | 0.0246 | 1 | 29 |
| 3 | 1575141 | 50117 | 0.0318 | 5 | 6 |
| 4 | 1084805 | 30247 | 0.0279 | 2 | 0 |
| 5 | 1814975 | 54199 | 0.0299 | 6 | 0 |
| 6 | 1422463 | 45598 | 0.0321 | 3 | 0 |
| 7 | 1399503 | 42668 | 0.0305 | 7 | 0 |
| 8 | 1398693 | 42582 | 0.0304 | 1 | 0 |
| 9 | 1186808 | 37586 | 0.0317 | 1 | 12 |
| 10 | 1059964 | 29439 | 0.0278 | 5 | 1 |
| 11 | 1561994 | 47038 | 0.0301 | 4 | 0 |
| 12 | 774062 | 20441 | 0.0264 | 4 | 3 |
| 13 | 756744 | 20401 | 0.0270 | 3 | 1 |
| 14 | 926563 | 27207 | 0.0294 | 4 | 0 |

**Suppl. Figure 1.**
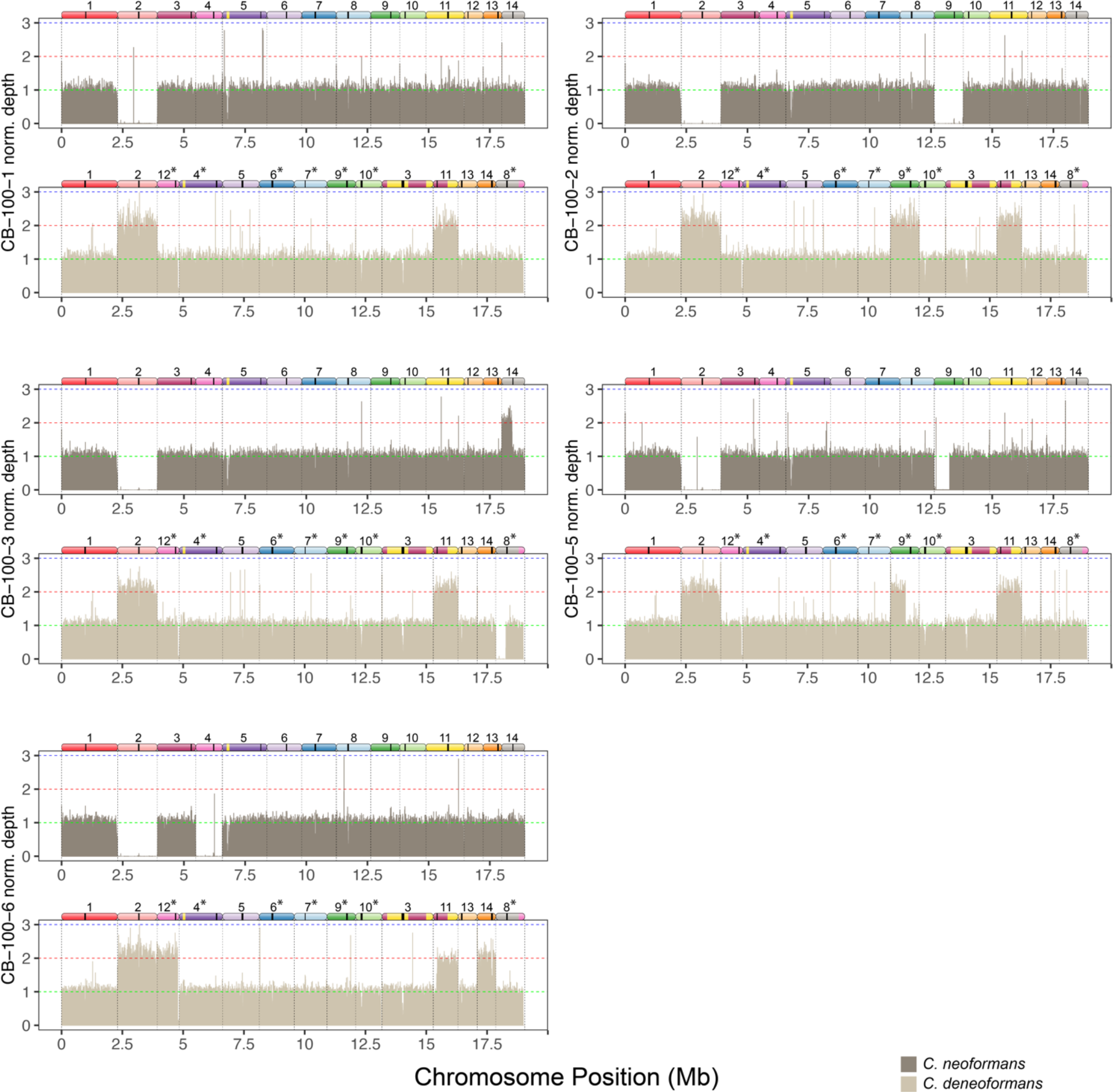
Mapping of reads to a hybrid reference genome of strains evolved in beer wort. Bars show mean depth in 5 kb intervals that have been normalized by dividing the depth of each interval by the mean coverage for each strain. For each strain the top panel shows mapping to the *C. neoformans* reference genome, and the bottom panel shows mapping to the *C. deneoformans* reference genome. Diagrams of homeologous chromosomes are shown above each reference genome where colors reflect homology and black stripes indicate centromeres. *C. deneoformans* chromosomes were rearranged to better reflect homology, and asterisks indicate chromosomes that have been reverse-complemented.

**Suppl. Figure 2.**
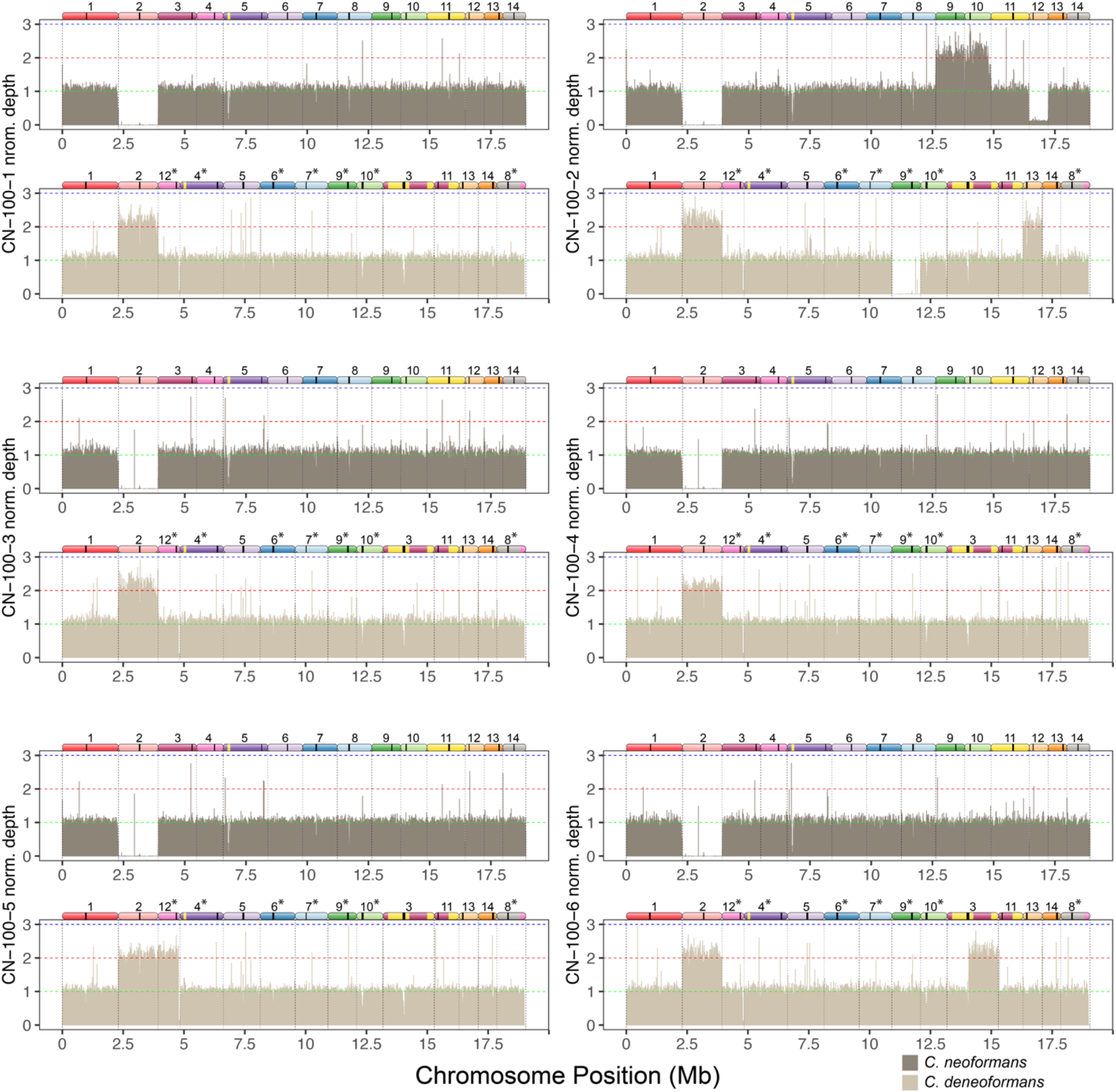
Mapping of reads to a hybrid reference genome of strains evolved in high salt. Bars show mean depth in 5 kb intervals that have been normalized by dividing the depth of each interval by the mean coverage for each strain. For each strain the top panel shows mapping to the *C. neoformans* reference genome, and the bottom panel shows mapping to the *C. deneoformans* reference genome. Diagrams of homeologous chromosomes are shown above each reference genome where colors reflect homology and black stripes indicate centromeres. *C. deneoformans* chromosomes were rearranged to better reflect homology, and asterisks indicate chromosomes that have been reverse-complemented.

**Suppl. Figure 3.**
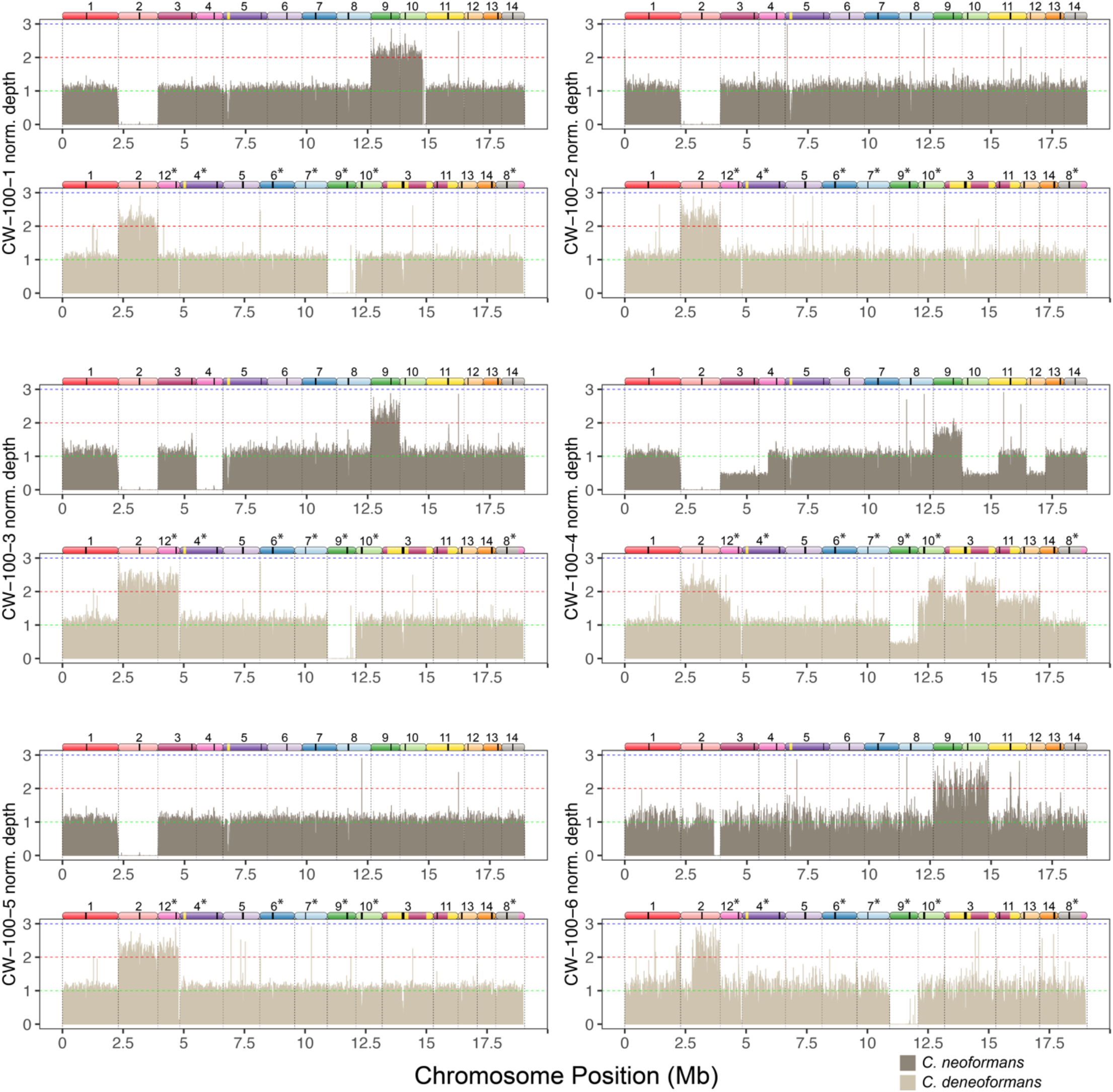
Mapping of reads to a hybrid reference genome of strains evolved in wine must. Bars show mean depth in 5 kb intervals that have been normalized by dividing the depth of each interval by the mean coverage for each strain. For each strain the top panel shows mapping to the *C. neoformans* reference genome, and the bottom panel shows mapping to the *C. deneoformans* reference genome. Diagrams of homeologous chromosomes are shown above each reference genome where colors reflect homology and black stripes indicate centromeres. *C. deneoformans* chromosomes were rearranged to better reflect homology, and asterisks indicate chromosomes that have been reverse-complemented.

**Suppl. Figure 4.**
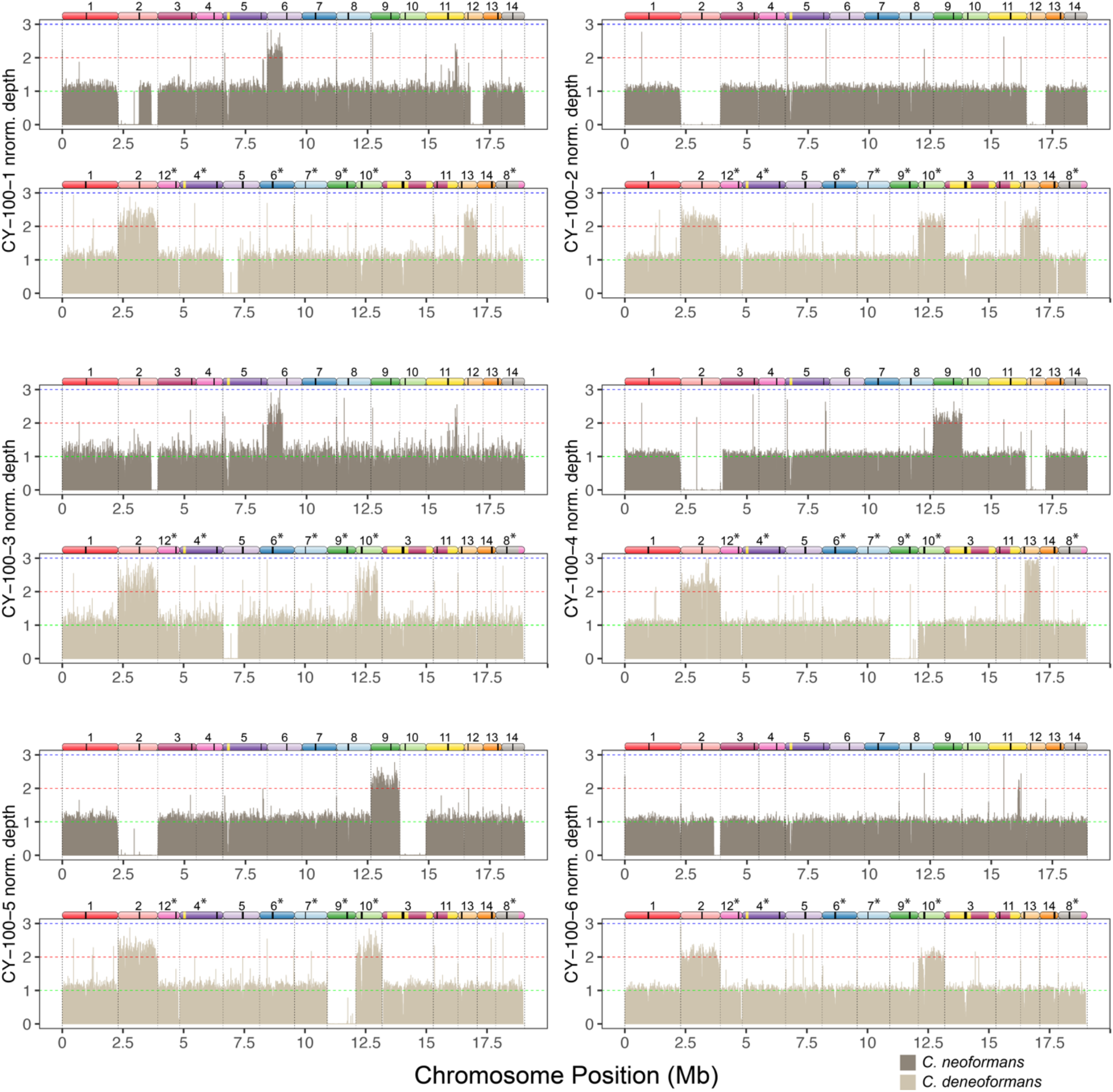
Mapping of reads to a hybrid reference genome of strains evolved in complete media (YPD). Bars show mean depth in 5 kb intervals that have been normalized by dividing the depth of each interval by the mean coverage for each strain. For each strain the top panel shows mapping to the *C. neoformans* reference genome, and the bottom panel shows mapping to the *C. deneoformans* reference genome. Diagrams of homeologous chromosomes are shown above each reference genome where colors reflect homology and black stripes indicate centromeres. *C. deneoformans* chromosomes were rearranged to better reflect homology, and asterisks indicate chromosomes that have been reverse-complemented.

**Suppl. Figure 5.**
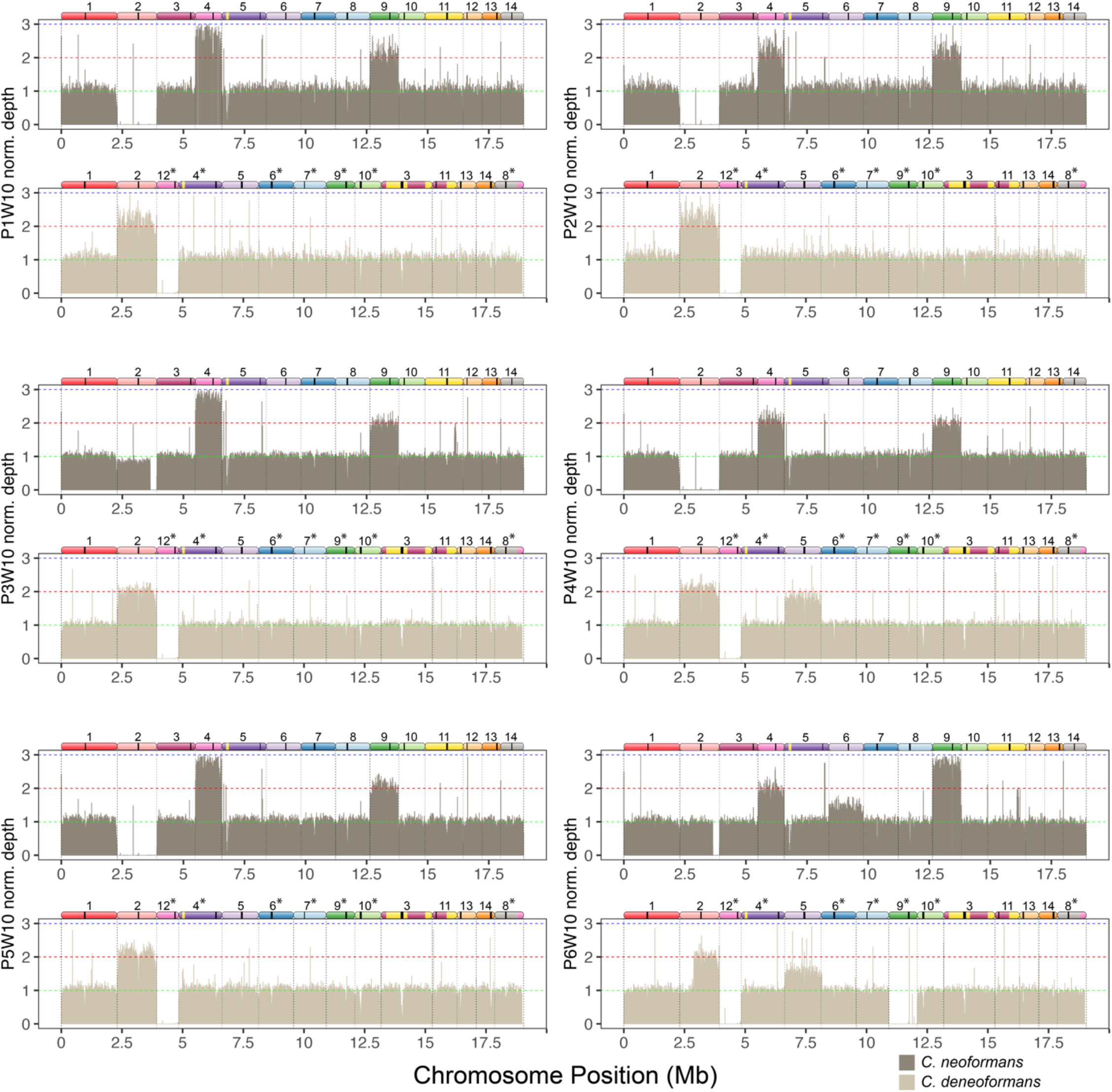
Mapping of reads to a hybrid reference genome of strains evolved in larvae. Bars show mean depth in 5 kb intervals that have been normalized by dividing the depth of each interval by the mean coverage for each strain. For each strain the top panel shows mapping to the *C. neoformans* reference genome, and the bottom panel shows mapping to the *C. deneoformans* reference genome. Diagrams of homeologous chromosomes are shown above each reference genome where colors reflect homology and black stripes indicate centromeres. *C. deneoformans* chromosomes were rearranged to better reflect homology, and asterisks indicate chromosomes that have been reverse-complemented.

**Suppl. Figure 6.**
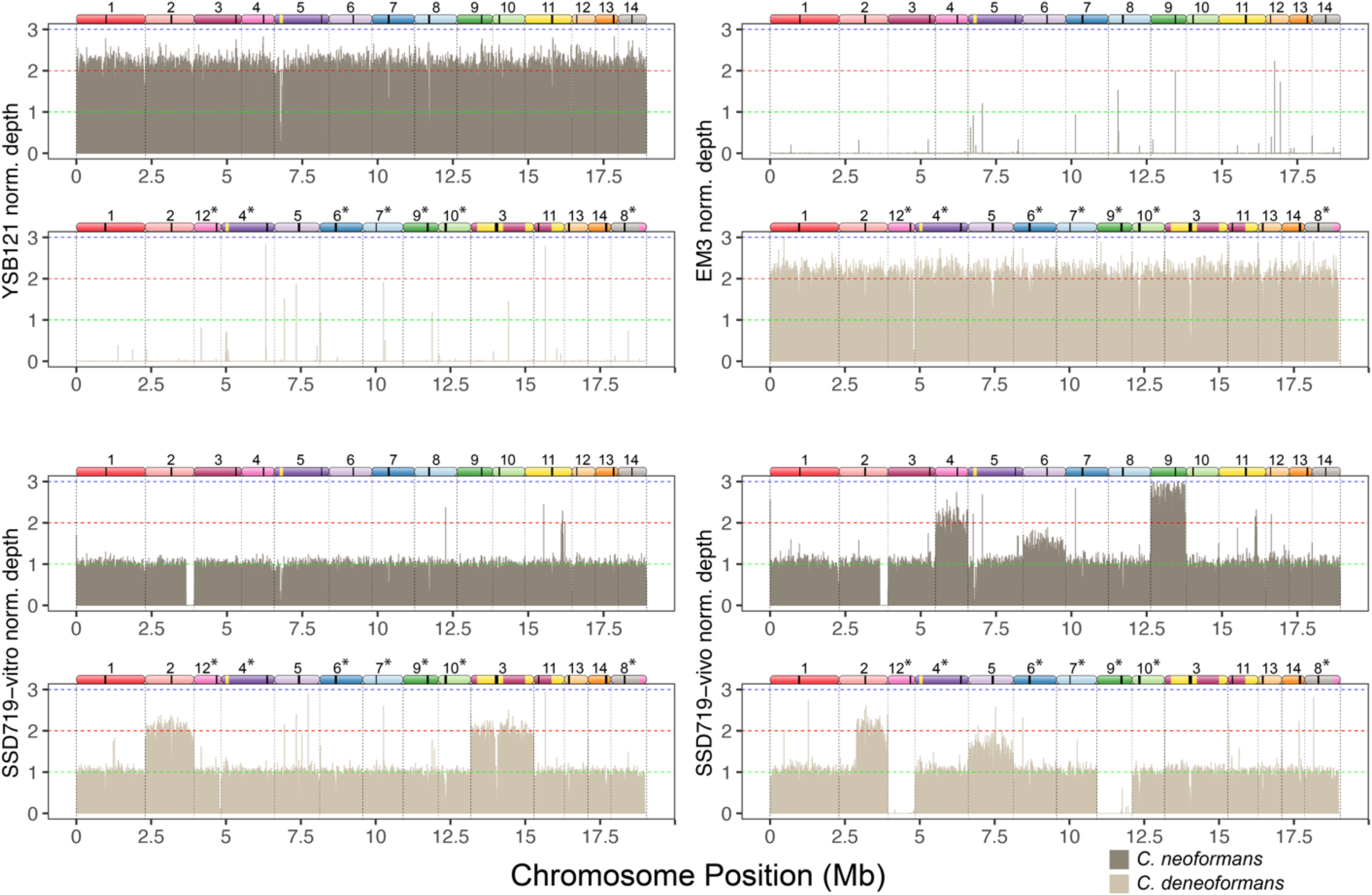
Mapping of reads to a hybrid reference genome of ancestral strains. Strains YSB121 and EM3 are haploid *C. neoformans* and *C. deneoformans* parents, respectively. Strain SSD719-in vitro was used to initiate the in vitro evolution populations and strain SSD719-in vivo was used to initiate the passages in larvae. Bars show mean depth in 5 kb intervals that have been normalized by dividing the depth of each interval by the mean coverage for each strain. For each strain the top panel shows mapping to the *C. neoformans* reference genome, and the bottom panel shows mapping to the *C. deneoformans* reference genome. Diagrams of homeologous chromosomes are shown above each reference genome where colors reflect homology and black stripes indicate centromeres. *C. deneoformans* chromosomes were rearranged to better reflect homology, and asterisks indicate chromosomes that have been reverse-complemented.

